# Chromatin-based mechanisms to coordinate convergent overlapping transcription

**DOI:** 10.1101/2020.03.27.011544

**Authors:** Soichi Inagaki, Mayumi Takahashi, Kazuya Takashima, Satoyo Oya, Tetsuji Kakutani

**Affiliations:** National Institute of Genetics, 1111 Yata, Mishima, Shizuoka, 411-8540, Japan; Department of Genetics, School of Life science, The Graduate University for Advanced Studies (SOKENDAI), Shonankokusaimura, Hayama, Kanagawa, 240-0193, Japan; Department of Biological Sciences, Faculty of Science, The University of Tokyo, 7-3-1 Hongo, Bunkyo-ku, Tokyo, 113-0033, Japan; PREST, Japan Science and Technology Agency, 4-1-8, Honcho, Kawaguchi, Saitama, 332-0012, Japan

## Abstract

In eukaryotic genomes, transcription units of genes often overlap with other protein-coding and/or noncoding transcription units^1,2^. In such intertwined genomes, coordinated transcription of nearby or overlapping genes would be important to ensure integrity of genome function; however, the mechanisms underlying this coordination are largely unknown^3-6^. Here, we show in *Arabidopsis thaliana* that genes with convergent orientation of transcription are major sources of overlapping bidirectional transcripts and that these bidirectionally transcribed genes are regulated by a putative LSD1 family histone demethylase, FLD^7,8^. Our genome-wide chromatin profiling revealed that FLD downregulated histone H3K4me1 in regions with convergent overlapping transcription. FLD localizes to actively transcribed genes where it colocalizes with elongating RNA polymerase II phosphorylated at Ser-2 or Ser-5 sites. Genome-wide transcription analyses suggest that FLD-mediated H3K4me1 removal negatively regulates bidirectional transcription by retaining the elongating transcription machinery. Furthermore, this effect of FLD on transcription dynamics is mediated by DNA topoisomerase I. Our study has revealed chromatin-based mechanisms to cope with overlapping bidirectional transcription, likely by modulating DNA topology. This global mechanism to cope with bidirectional transcription could be co-opted for specific epigenetic processes, such as cellular memory of responses to environment^9^.

## Main text

Methylation of histone H3 lysine-4 (H3K4me) is associated with active gene transcription. While H3K4 trimethylation (H3K4me3) occurs around transcription start sites (TSSs), H3K4 dimethylation (H3K4me2) and monomethylation (H3K4me1) occur in downstream regions (bodies), where transcription elongation occurs. The role of H3K4me1 in enhancers has been well studied in metazoans^10^, but H3K4me1 more generally occurs in gene bodies among eukaryotes, including yeasts and plants. Our previous genetic studies in a model plant *Arabidopsis thaliana* (hereafter Arabidopsis) showed the importance of H3K4me1 in the gene body in regulating active and inactive chromatin states^11^. In this report, by using genetic and genomic approaches, we uncover a novel mechanism underlying the global regulation of bidirectional transcription by controlling H3K4me1 in the gene body.

### FLD decreases H3K4me1 around TTS of convergently transcribed genes

FLOWERING LOCUS D (FLD) is one of the four Arabidopsis orthologs of human LSD1 (Lysine-Specific Demethylase 1), a demethylase of H3K4me2 and H3K4me1^12^. FLD has been shown to regulate flowering, the transition from vegetative growth to reproductive development^7^. Although it has been proposed that FLD silences the key flowering-controlling gene, *FLOWERING LOCUS C* (*FLC*), through removal of H3K4me2 there^8,13^, the genome-wide function of FLD remains unexplored. In contrast to the results of previous reports, our chromatin immunoprecipitation coupled with high-throughput sequencing (ChIP-seq) results showed that H3K4me1, rather than H3K4me2, significantly increased within the body of *FLC* gene in *fld* mutant compared with wild-type (WT) plants (Fig. 1a, b, Extended Data Figs. 1a, 2a). In addition, many other genes exhibited increased levels of H3K4me1 in *fld* (Fig. 1a). The increase in H3K4me1 in gene bodies was predominantly observed towards the transcriptional termination site (TTS; Fig. 1c, Extended Data Fig. 2b, c). Notably, the increase in H3K4me1 is most obvious when the downstream gene was convergently transcribed in relation to the target gene (Fig. 1b, d, Extended Data Fig. 1b, c). Based on the orientation of their downstream genes, genes can be classified as convergent or tandem. Within the whole Arabidopsis genome, approximately half (15,491/32,355) of the genes are convergent, and the other half are tandem; however, most of the genes exhibiting increased H3K4me1 in *fld* are convergent (Fig. 1d). The distance between the TTS of convergently transcribed genes correlates with the response of H3K4me1 to the *fld* mutation; H3K4me1 increased more when the downstream genes were close to or partly overlapped with the target genes (Fig. 1e, Extended Data Fig. 1d). These results demonstrate that FLD decreases H3K4me1 around the TTS of convergently transcribed genes.

**Fig. 1.**
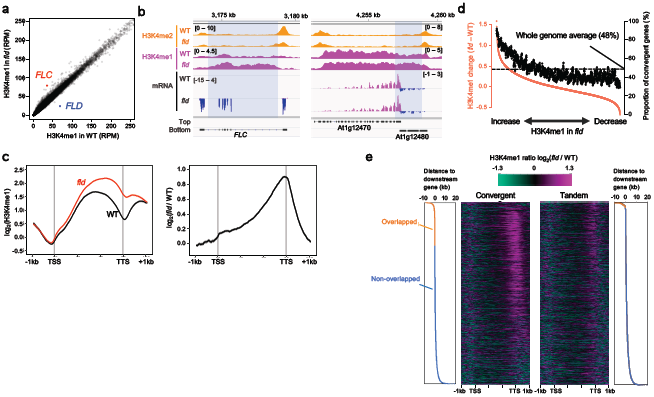
FLD, LD, and SDG26 decrease H3K4me1 around TTS of convergently transcribed genes. **a**, Scatter plot comparing H3K4me1 between WT and *fld*. Each dot represents the read counts per million mapped reads (RPM) within each gene. *FLC* accumulating H3K4me1 and *FLD* disrupted in *fld* are highlighted in red and blue, respectively. **b**, Patterns of H3K4me2, H3K4me1, and mRNA (top strand, magenta; bottom strand, dark blue) around the *FLC* locus (left) and a pair of convergent genes (right). Normalized coverage (per million mapped reads) is shown. The numbers in parentheses indicate the range of values. Gene structures (box, exon; line, intron) in the top and bottom strands are shown. The shaded areas have increased H3K4me1 levels in *fld*. **c**, Averaged profiles of H3K4me1 in WT and *fld* (left) and H3K4me1 change (right) around genes with increased H3K4me1 in *fld* (1842 genes). The ribbon indicates SEM. **d**, Sliding window analysis (window size, 150; sliding size, 20) showing the proportion of convergent genes in relation to H3K4me1 changes between WT and *fld*. All genes were ranked and grouped by the difference in H3K4me1 levels between WT and *fld* (red plots and left axis) and the proportion of convergent genes in each window is shown by a black plot and the right axis. The dashed line indicates the proportion of convergent genes within the whole genome (48%). **e**, Heatmaps showing H3K4me1 changes between WT and *fld* around genes sorted by the distance to downstream genes for convergent and tandem genes separately. The distance (plus value in blue, nonoverlapping; minus value in orange, overlapping) to the downstream gene for each gene is plotted in graphs on both sides.

To determine whether the presence of a convergently transcribed gene is a prerequisite for FLD to remove H3K4me1 from the gene body, we chose a pair of convergent genes, one of which (At1g77120) exhibits increased H3K4me1 in *fld* (Extended Data Fig. 3a) and questioned whether disruption of the downstream gene (At1g77122) by transfer DNA (T-DNA) insertion affects H3K4me1 levels in the body of At1g77120. Two independent T-DNA insertions blocked the transcription of At1g77122 and increased H3K4me1 in the At1g77120 body (Extended Data Fig. 3a), supporting the idea that convergent transcription is a trigger for H3K4me1 demethylation by FLD.

### FLD localizes to TTS of actively transcribed genes

We then analyzed the genome-wide localization of the FLD protein by the use of transgenic plants expressing FLAG-tagged FLD proteins, which phenotypically complemented the *fld* mutant. Enrichment of FLD binding was observed around the TTS of the genes that increased H3K4me1 in *fld* (Fig. 2a, Extended Data Fig. 4). The enrichment of FLD proteins around the TTS was positively correlated with the expression levels in WT plants (Fig. 2b). We screened for chromatin features that predict FLD localization via random forest analysis (Fig. 2c). Among the chromatin and genomic features analyzed, the best predictors were phosphorylation of Ser-2 and Ser-5 (Ser2P and Ser5P, respectively) of the carboxy-terminal domain (CTD) of RNA polymerase II (RNAPII), which are implicated in multiple steps of transcription-coupled processes, such as elongation and termination^14^, around the TTS (Fig. 2c). Indeed, the levels of RNAPII-Ser2P and RNAPII-Ser5P around the TTS by themselves could predict the FLD localization with high accuracy (Extended Data Fig. 5a), and the genome-wide profiles and intragenic patterns of FLD localization resemble those of RNAPII-Ser2P and RNAPII-Ser5P (Extended Data Fig. 5b-e). Consistent with the idea that the transcription of target genes is important for FLD recruitment, we found that FLD localization to the TTS of *FLC* was induced in the plants actively expressing *FRIGIDA* (*FRI*), which upregulates transcription of *FLC*^15,16^ (Fig. 2d, Extended Data Fig. 6). Together, these results suggest that FLD is recruited preferentially to the TTS of actively transcribed genes, colocalizing with the elongating RNAPII.

**Fig. 2.**
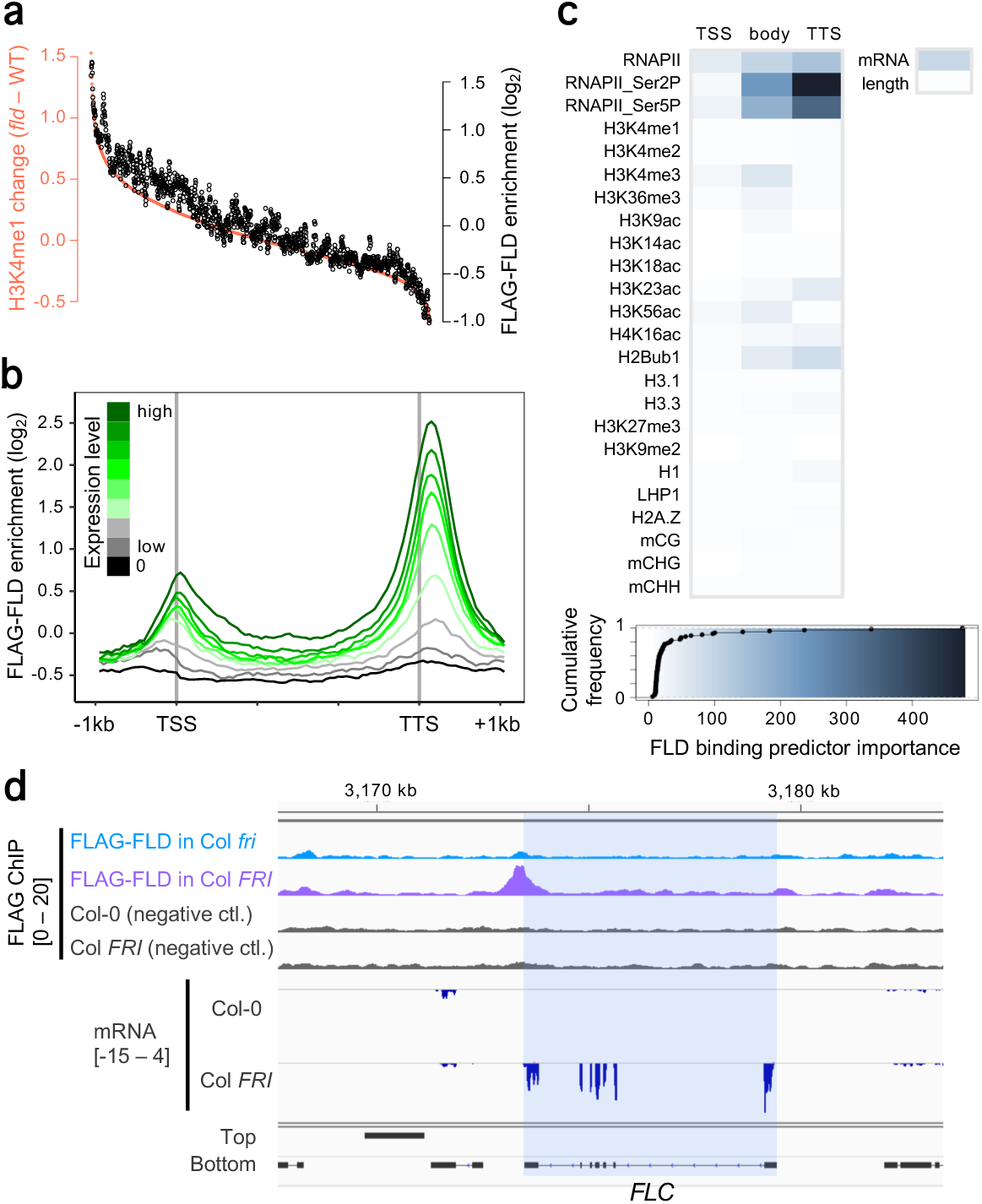
FLD localizes to TTS of actively transcribed genes. **a**, Sliding window analysis showing FLAG-FLD enrichment level in relation to H3K4me1 changes between WT and *fld*. All the genes were grouped like in Fig. 1d, and FLAG-FLD enrichment relative to the negative control is shown by black plots and the right axis. **b**, FLAG-FLD enrichment pattern around genes grouped according to expression level in WT. **c**, Relative importance of chromatin features for predicting FLD binding to each gene, analyzed via random forest. The chromatin features around TSS, TTS, and gene body were analyzed separately. The bottom panel shows the color code for the relative importance value and cumulative frequency of value for each feature. **d**, Patterns of FLAG-FLD enrichment and mRNA around *FLC* (shaded), comparing Col *fri* and Col *FRI*.

### FLD controls transcription dynamics of bidirectionally transcribed genes

Our genome-wide analyses revealed that convergently transcribed genes are preferred targets of FLD. However, the *FLC* locus is an interesting exception; although H3K4me1 increases throughout the gene body of *FLC* in *fld*, the annotated gene downstream of *FLC* is not convergent with respect to *FLC* (Extended Data Fig. 1a). Notably, however, several noncoding RNA species have been reported to be present at the *FLC* locus. A set of antisense noncoding RNAs, *COOLAIR*, plays a key role to trigger transcriptional silencing of *FLC* in response to low ambient temperature^17,18^. To determine whether antisense transcription is globally associated with FLD targets, we analyzed the genome-wide transcriptome using mRNA sequencing (mRNA-seq) and chromatin-bound RNA sequencing (chrRNA-seq). While mRNA represents mostly the soluble mature fraction of RNA in cells, chrRNA is enriched for the nascent pre-mRNA associated with RNAPII on chromatin before or during splicing (Extended Data Fig. 7a). Remarkably, the chrRNA-seq results showed that many genes with highly increased H3K4me1 in *fld* have high levels of antisense transcripts, which are mostly undetectable by mRNA-seq (Fig. 3a, b, Extended Data Figs. 7b, c, 8a). These results suggest that FLD function is linked to overlapping bidirectional transcription.

**Fig. 3.**
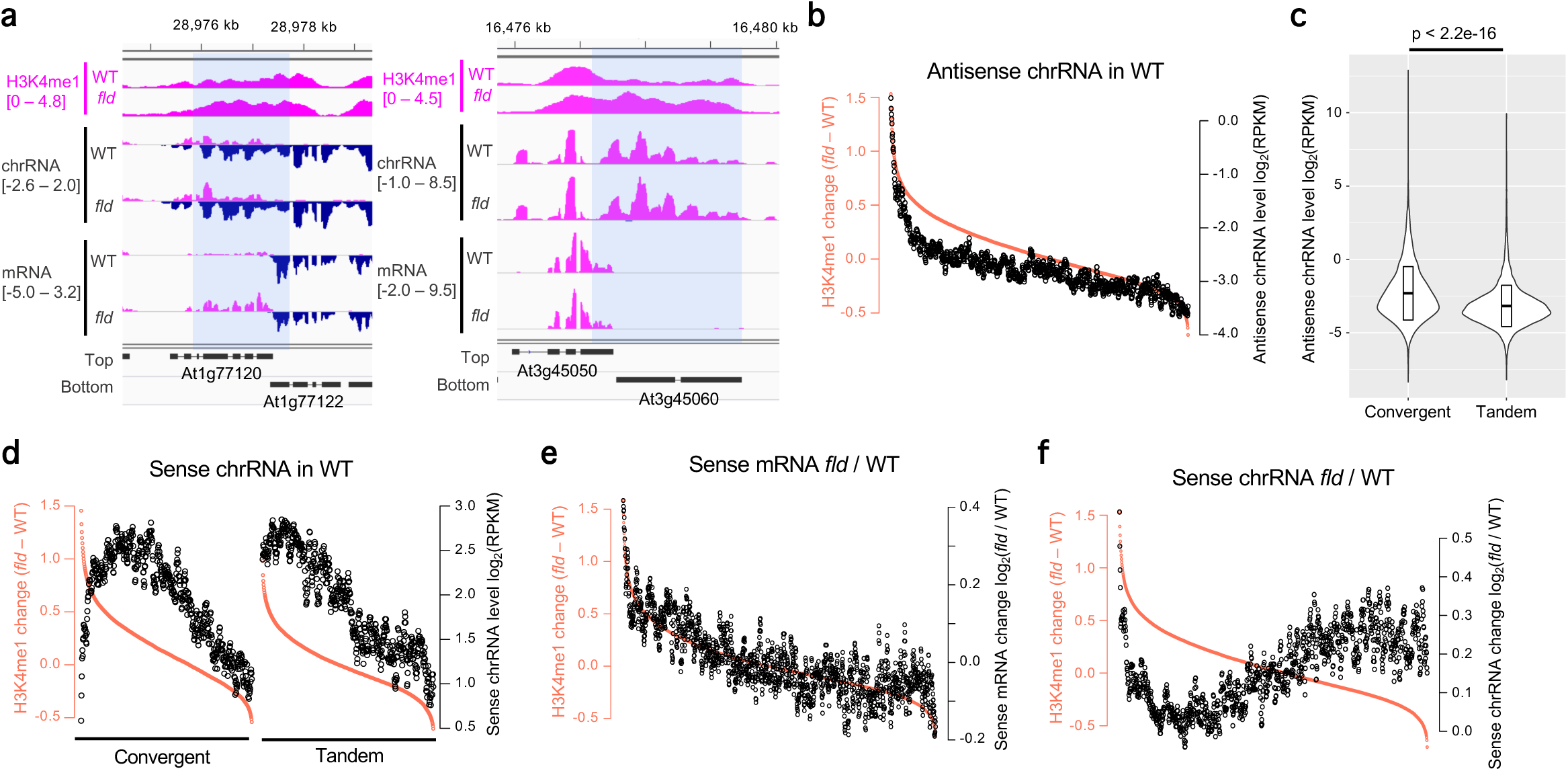
FLD controls transcription dynamics of bidirectionally transcribed genes. **a**, Patterns of H3K4me1, chrRNA, and mRNA of convergent genes with antisense chrRNA. The shaded areas have increased H3K4me1 in *fld* compared to WT. **b**, Sliding window analysis showing the relationship of changes in H3K4me1 between WT and *fld* to antisense chrRNA levels in WT. All the genes were grouped like in Fig. 1d, and the antisense chrRNA levels in WT are shown by black plots and the right axis. RPKM, read counts per kilobases per million mapped reads. **c**, Violin plots of antisense chrRNA levels in convergent and tandem genes. The boxes inside represent the means±SDs. The *p*-value is based on Welch’s *t*-test comparing two groups of genes. **d**, Sliding window analysis showing the relationship of changes in H3K4me1 levels between WT and *fld* to sense chrRNA levels in WT, shown separately for convergent and tandem genes. **e, f**, Sliding window analysis showing the relationship of changes in H3K4me1 to changes in mRNA level (**e**) or to changes in chrRNA levels (**f**) between WT and *fld*.

Interestingly, the antisense transcripts detected by chrRNA-seq were more enriched in convergent genes than in tandem genes (Fig. 3c, Extended Data Fig. 8b), suggesting that, at least in the Arabidopsis genome, convergent transcription is a major source of antisense transcripts, likely reflecting readthrough transcription of downstream genes (Fig. 3a). Consistent with the idea of readthrough being a contributing factor, high levels of sense transcripts were detected in genes downstream of the genes with enriched antisense transcripts, especially in convergent cases (Extended Data Fig. 8c). Importantly, although sense and antisense chrRNA of the same gene were generally positively correlated, convergent genes with the highest amount of antisense chrRNA were relatively repressed (Extended Data Fig. 8d), implying that antisense transcription and/or transcripts could have a negative impact on sense transcription.

FLD removes H3K4me1 from genes that have antisense transcripts, which may be associated with gene silencing. Indeed, sense chrRNA is low in the convergent genes from which FLD removes H3K4me1 most strongly (Fig. 3d, left). Furthermore, our analyses of chrRNA and mRNA in the *fld* mutant revealed impacts of FLD on transcription dynamics. Generally, *fld* increased the level of sense mRNA in genes with increased H3K4me1 (Fig. 3e), suggesting that increased H3K4me1 is associated with transcriptional activation. Sense chrRNA in genes with highly increased H3K4me1, including *FLC*, also increased in *fld* (Fig. 3f, Extended Data Fig. 7a), which is consistent with a previous proposal that *FLC* is transcriptionally upregulated not only at elongation but also at initiation level in *fld*^13^.

Interestingly, however, sense chrRNA in genes with a moderate increase in H3K4me1 decreased rather than increased in *fld* (Fig. 3f). These results suggest that the increase in H3K4me1 in *fld* accelerates the release of mRNA from chromatin, most likely by increasing the elongation rate. In addition to the effect on the dynamics of sense transcription, *fld* also has differential effects on antisense chrRNA and mRNA (Extended Data Fig. 8e); in genes with a strong increase in H3K4me1 in *fld*, antisense chrRNA is low, while antisense mRNA is high, suggesting that FLD functions to retain the elongating transcription machinery not only for sense transcription but also for antisense transcription. Our hypothesis that FLD regulates transcription elongation was also supported by our RNAPII ChIP-seq analyses, the results of which showed that *fld* induces a decrease in RNAPII levels around the TTS of genes that have elevated levels of H3K4me1 (Fig. 4a, Extended Data Fig. 9a-d). Globally, the effects of *fld* on H3K4me1 and RNAPII around the TTS were negatively correlated (Fig. 4b). These results are consistent with the idea that H3K4me1 facilitates the passage of transcription machinery and that FLD counteracts this process through removal of H3K4me1 around the TTS. In contrast, the effects of *fld* on RNAPII levels around the TSS resembled those on chrRNA (Figs. 3f, 4c), reflecting the effect of H3K4me1 on both transcription initiation and elongation when H3K4me1 changes are relatively large.

**Fig. 4.**
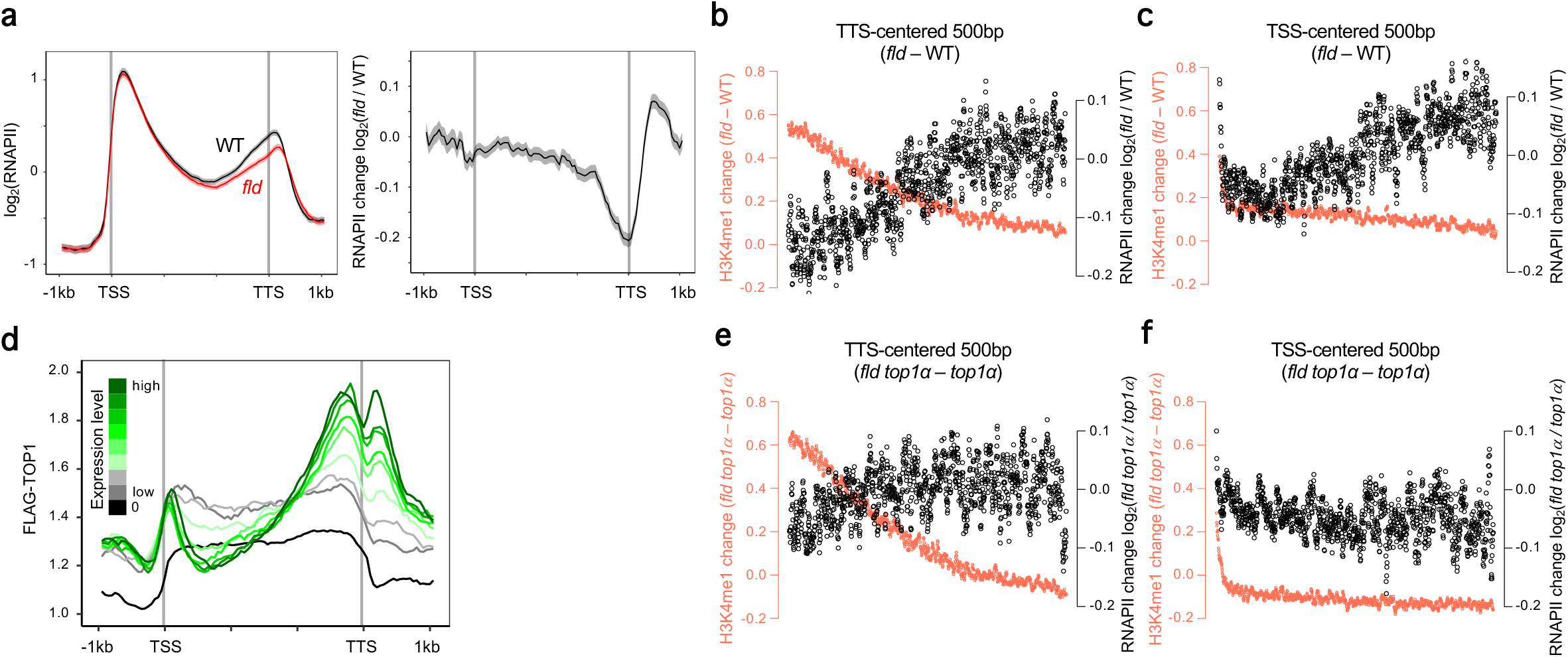
FLD controls transcription dynamics around TTS through DNA topoisomerase I function. **a**, Averaged profiles of RNAPII in WT and *fld* (left) and RNAPII changes (right) around genes with increased H3K4me1 in *fld* (1842 genes). The ribbon indicates SEM. **b, c**, Sliding window analysis showing the relationship of changes in H3K4me1 to changes in RNAPII levels around TTS (**b**) and TSS (**c**) between WT and *fld*. All the genes were grouped like those in Fig. 1d, and H3K4me1 changes in the TTS- or TSS-centered 500 bp area are shown by red plots and the left axis. The ratio of RNAPII levels in the TTS- or TSS-centered 500 bp area between WT and *fld* is shown by black plots and the right axis. **d**, FLAG-TOP1 localization pattern around genes grouped according to their expression levels in WT. **e, f**, Sliding window plot for comparisons of H3K4me1 and RNAPII changes around TTS (**e**) and TSS (**f**) between *top1α* and *fld top1α*.

### FLD controls transcription dynamics around TTS through DNA topoisomerase I function

Our results suggest that the plausible basis of the FLD-regulated transcription dynamics is by the modulation of transcription elongation. Transcription elongation would have additional impacts, especially on convergently transcribed genes. Transcription is known to impose positive DNA supercoiling ahead of the polymerase and negative DNA supercoiling behind the polymerase^19,20^. Changes in chromatin modification and transcription dynamics within convergent and bidirectional genes in *fld* led us to ask if the regulation occurs to cope with superhelicity or torsional stress, because strong positive supercoils would accumulate between convergent gene pairs or within overlapping regions of those pairs^21^. Type I and type II DNA topoisomerases make transient single-stranded and double-stranded breaks, respectively, to alter superhelicity and relax the supercoils imposed by DNA replication or transcription^20^. It has recently been reported that in human cells, TOP1 functions within the transcriptional machinery of RNAPII and helps RNAPII travel along the gene body by altering the DNA topology ahead of RNAPII^22^. Indeed, our ChIP-seq results for a DNA topoisomerase I protein in Arabidopsis, TOP1α, show that TOP1α localizes to active genes towards the TTS (Fig. 4d). Furthermore, the loss-of-function *top1α* mutant reportedly exhibits an early-flowering phenotype, and the *top1α* mutation partly suppress the late-flowering phenotype of the *fld* mutant through downregulation of *FLC*^23^. We therefore tested whether *top1α* also suppresses the effect of *fld* on transcription dynamics globally. Our RNAPII ChIP-seq results for *top1α* and *fld top1α* double mutant revealed that the effects of *fld* on RNAPII levels around the TTS and TSS were suppressed in the *top1α* mutant background (Fig. 4e, f), suggesting that FLD slows transcription elongation via modulation of DNA topology through TOP1α activity.

## Discussion

Although previous reports in yeast and humans have characterized divergent transcription events from bidirectional promoters as sources of antisense transcripts^3,24,25^, impacts of convergent transcription are less understood. Our chrRNA analyses revealed that convergent transcription events are major sources of antisense transcripts and are associated with gene silencing (Fig. 3a, c, d). In addition, the transcription dynamics of genes with antisense transcripts are coordinated by FLD and associated factors, most likely through removal of H3K4me1. One plausible scenario is that H3K4me1 in the gene body accelerates transcription elongation, possibly through modulating DNA topology, and that is antagonized by FLD to decelerate elongation around the TTS, especially when downstream or overlapping genes are convergently transcribed (Extended Data Fig. 10).

Importantly, the removal of H3K4me1 from the gene body affects not only transcription elongation but also transcription initiation (Fig. 3e, f). Consistent with this observation, the results of our previous study showed that the loss of H3K4me1 in the gene body mediates constitutive gene silencing triggered by H3K9me^11^. It would be interesting to investigate whether changes in transcription elongation and DNA topology generally mediate the effects of H3K4me1 in the gene body on transcription initiation.

The mechanism we uncovered here is co-opted by plants for epigenetic silencing of *FLC*, which is triggered by cold in the winter and controls developmental phase transition via antisense transcription^9,16-18^. Overlapping antisense transcripts are also found in other epigenetic systems, such as in mammalian imprinted genes and in the X-chromosome inactivation center^26,27^. It would be interesting to learn how gene body chromatin modifications, torsional stress and transcription dynamics interact in these and other systems.

## Acknowledgements

We thank Akiko Terui for technical assistance, and Taku Takahashi for plant material. This work used the Vincent J. Coates Genomics Sequencing Laboratory at UC Berkeley, supported by NIH S10 OD018174 Instrumentation Grant. Computations were partially performed on the NIG supercomputer at NIG, Japan. This work was supported by grants from JST PREST (JPMJPR17Q1) to SI, and JST CREST (JPMJCR15O1) and JSPS (26221105, 15H05963 and 19H00995) to TK.

## Author contributions

SI and TK designed the study. SI, MT, and KT performed experiments. SI analyzed the data. SO conducted random forest analysis. SI and TK wrote the manuscript with incorporating comments from other authors.

## Methods

### Plant materials and growth condition

*Arabidopsis thaliana* strain Columbia-0 (Col-0) was used as “wild type (WT)”. The *fld-4* and *top1α-1* mutants were characterized^7, 31^. Plants were grown on Murashige and Skoog (MS) media supplemented with 1% sucrose and solidified with Bacto agar (Difco) in long day condition (16/8-h photoperiod).

### Plasmid construction and plant transformation

For pFLD::3xFLAG-FLD-HA, a genomic region spanning the promoter was amplified with following primers; 5’-AATTCTAGTTGGAATGGGTTATGCTGGCGAACTCACTCC-3’ and 5’-CTATATCGTGATCTTTGTAATCTCCATCGTGATCTTTGTAATCCATCTGCTCAAAACTAGGGTTAGAG-3’, and coding region with intron until just before its stop codon was amplified with following primers; 5’-GAGATTACAAAGATCACGATATAGATTACAAAGATGATGATGATAAGGTCTCATTCTCCGCACCAAAG-3’ and 5’-GATCCTTATGGAGTTGGGTTTTAAGCGTAATCTGGAACATCGTATGGGTATTGTTCAATCTTTTTCATCGTCTCAC-3’. The amplified fragments were assembled into HpaI-linearlized pPLV01 vector^32^ by using the NEBuilder HiFi DNA Assembly Master Mix (NEB) and cloned in *E. coli*. pTOP1α::3xFLAG-TOP1α was similarly made with following primers; 5’-TATCGAATTCTAGTTGGAATGGGTTGATATTTAACAAGTCTCTTCTGG-3’ and 5’-CTATATCGTGATCTTTGTAATCTCCATCGTGATCTTTGTAATCCATTCCCGAAAGAACAACGTTG-3’ for the promoter, and 5’-GAGATTACAAAGATCACGATATAGATTACAAAGATGATGATGATAAGGGCACTGAAACAGTTTCAAAAC-3’ and 5’-GATCGGATCCTTATGGAGTTGGGTTTGTTTTTGTTCTTCTTCGGATTTTC-3’ for the gene body. Plasmid from a single *E. coli* colony was checked with sequencing and was transformed into *Agrobacterium tumefaciens* GV3101::pMP90 containing pSOUP^33^ using electroporation. Heterozygous *fld-4*/+ or *top1α*/+ plants were transformed with the plasmids using the floral dip method, and the transgenic lines having homozygous transgenes and homozygous for *fld-4 or top1α* mutation, respectively, were selected in the later generation.

### ChIP-seq

Chromatin immunoprecipitation (ChIP) for histone modifications was carried out as described in ref. 11. Approximately 1.5 gram of 14-day-old whole seedlings grown on MS media was frozen with liquid nitrogen, ground into fine powder, cross-linked with 1% formaldehyde, and nuclear-extracted according to ref. 34. Nuclear pellet was lysed with 150 µl of lysis buffer (50 mM Tris-HCl, pH 7.8, 10 mM EDTA, 1% SDS), diluted with 800µl of dilution buffer (50 mM Tris-HCl, pH 7.8, 0.167 M NaCl, 1.1 % Triton X-100, 0.11% Sodium Deoxycholate), and sonicated using a Covaris S2 Focused-ultrasonicator (Covaris) and milliTUBE 1 ml AFA Fiber (Covaris) with the following settings: power mode, frequency sweeping; time, 20 min; duty cycle, 5%; intensity, 4; cycles per burst, 200; and temperature (water bath), 4–6°C. The sonicated chromatin was then centrifuged at 13,000 g for 3 min and supernatant was transferred to a new 1.7 ml tube. Immunoprecipitation was performed by incubating diluted chromatin solution with rabbit anti-H3K4me1 (ab8895; Abcam), rabbit anti-H3K4me2 (ab32356; Abcam), or rabbit anti-H3 (ab1791; Abcam) overnight at 4°C, and incubating for 2 hours with Dynabeads Protein G (Veritas). The incubated beads were washed once with 1 ml of low salt wash buffer (50 mM Tris-HCl, pH 7.8, 150 mM NaCl, 1mM EDTA, 0.1% SDS, 1% Triton X-100, 0.1% Sodium Deoxycholate, cOmplete EDTA-free Protease Inhibitor Cocktail (Sigma-Aldrich)), twice with 1 ml of high salt wash buffer (wash buffer with 500 mM NaCl), once with 1 ml of LiCl buffer (10 mM Tris-HCl, pH 7.8, 1 mM EDTA, 0.25 M LiCl, 1% IGEPAL CA-630, 1% Sodium Deoxycholate), once with 1 ml of TE buffer (10 mM Tris-HCl, pH 7.8, 1 mM EDTA). DNA was eluted from the beads and reverse-crosslinked by adding 100 µl elution buffer (10 mM Tris-HCl, pH 7.8, 0.3 M NaCl, 5 mM EDTA, 0.5 % SDS) and incubating overnight at 65 °C. The DNA samples were then treated with RNase A (Nippon Gene) for 30 min and with Proteinase K (ThermoFisher) for 2 h at 37°C, and purified using the Monarch PCR & DNA Cleanup Kit (NEB). Collected DNA was quantified with the Qubit dsDNA High Sensitivity Assay kit (ThermoFisher). ChIP-seq library was made using the ThruPLEX DNA-seq Kit (Takara), and dual size-selected using Agencourt AMPure XP (Beckman Coulter) to enrich 200 – 500-bp fragments. The libraries were pooled and 50-bp single-end sequences were obtained using the HiSeq4000 sequencer (Illumina) in Vincent J. Coates Genomics Sequencing Laboratory at UC Berkeley. Two independent biological replicates were analyzed for each experiment except for the experiment in Extended Fig. 3b, which was performed once.

Genome-wide localization pattern of FLD and TOP1α proteins were analyzed by using 3xFLAG-FLD-HA transgenic plants in the *fld* mutant background and 3xFLAG-TOP1α transgenic plants in the *top1α* mutant background, respectively, and non-transgenic control (WT Col-0). Approximately 1.5 gram of 14-day-old whole seedlings grown on MS media was frozen with liquid nitrogen, ground into fine powder, cross-linked with 1% formaldehyde and EGS (ethylene glycol bis(succinimidyl succinate); ThermoFisher), and nuclear-extracted as the histone modification ChIP. ChIP was performed similar as the histone modification ChIP described above with the following modifications. Nuclear pellet was lysed with 1 ml of low salt wash buffer and used for sonication. 1 µg of mouse anti-FLAG M2 monoclonal antibody (Sigma-Aldrich) was used. Washed twice with low salt wash buffer, twice with medium salt wash buffer (300 mM NaCl), and once with TE buffer. Two biological replicates were analyzed for each experiment, except for the experiment in Col *FRI* (Fig. 4d and Extended Data Fig. 6), which was performed once.

RNAPII ChIP-seq was performed as the histone modification ChIP-seq with the following modifications. Samples were cross-linked with 0.5% formaldehyde. Nuclear pellet was lysed with low salt wash buffer and sonicated with the following settings: power mode, frequency sweeping; time, 15 min; duty cycle, 5%; intensity, 3; cycles per burst, 200; and temperature (water bath), 4–6°C. 1 µg of mouse anti-RNA Polymerase II CTD monoclonal antibody (CMA601)^35^ or Monoclonal Pol II Antibody – ChIP-seq grade (Diagenode) for RNAPII, mouse anti-Phospho RNA Polymerase II CTD (Ser2) monoclonal antibody (CMA602)^35^ for RNAPII Ser2P, or mouse anti-Phospho RNA Polymerase II CTD (Ser5) monoclonal antibody (CMA603)^35^ for RNAPII Ser5P were used. Washed once with low salt wash buffer, once with high salt wash buffer, once with half-detergent LiCl buffer (0.5% IGEPAL CA-630, 0.5% Sodium Deoxycholate) and once with TE buffer. Two biological replicates were analyzed for each experiment.

### mRNA-seq

Total RNA was isolated from 14-day-old seedlings grown on MS media, using the RNeasy Plant Mini Kit (Qiagen), and treated with DNase I (Takara). Libraries for mRNA-seq were constructed using the KAPA Stranded mRNA-seq Kit according to the manufacturer’s instruction. 50-bp single-end sequences were obtained using the HiSeq4000 sequencer (Illumina) in Vincent J. Coates Genomics Sequencing Laboratory at UC Berkeley. Three biological replicates were analyzed for each experiment, except for mRNA-seq in Col *FRI* (Fig. 4d and Extended Data Fig. 6) and mRNA-seq in T-DNA lines of At1g77120-At1g77122 (Extended Data Fig. 3a), which were performed once.

### chrRNA-seq

First, nuclei were extracted from approximately 2 grams of 14-day-old whole seedlings grown on MS media following the protocol in ref. 36 with the following modifications. Formaldehyde crosslinking was omitted, 25 µM of α-amanitin (Sigma-Aldrich) was added throughout the experiment to stop transcription, 0.08U/µl of RNase Inhibitor (Toyobo) was added throughout the experiment, and nylon filters (100 µm and 40 µm pore sizes sequentially) instead of Miracloth were used to remove debris. Then, nuclei were lysed to remove nucleoplasmic fraction through the following process. Isolated nuclei were resuspended in 200 µl of NUN1 buffer (20 mM Tri-HCl, pH 8.0, 75mM NaCl, 0.5 mM EDTA, 50% Glycerol and 1x cOmplete EDTA-free Protease Inhibitor Cocktail) followed by 1 ml NUN2 buffer (20 mM HEPES-KOH pH 7.6, 7.5 mM MgCl_2_, 0.2 mM EDTA, 300 mM NaCl, 1 M Urea, 1% NP40, cOmplete EDTA-free Protease Inhibitor Cocktail). The solution was incubated at 4°C for 15 min, with vortexing every 3 min. The chromatin pellet was precipitated by performing 13,000 rpm centrifuge at 4°C for 10 min. The chromatin pellet was resuspended in 400 µl of lysis buffer (10 mM Tris-HCl, pH 7.8, 10 mM EDTA, 0.5 % SDS), and chrRNA was extracted using phenol-chloroform extraction and ethanol precipitation. At this point ∼3 µg of RNA was typically collected. The remaining DNA was depleted with 2 U of Baseline-ZERO DNase (Epicentre) and ribosomal RNA was depleted using the Ribo-Zero Plant rRNA removal kit (Epicentre). chrRNA-seq libraries were constructed from 50 ng of purified RNA using the KAPA Stranded mRNA-seq Kit, skipping poly-A selection step and performing RNA fragmentation step at 94°C for 4 min. 150-bp paired-read sequences were obtained using the HiSeq X Ten sequencer (Illumina) in Macrogen Japan Corp.

### Data analysis

For mRNA-seq, quality-filtered reads were mapped onto the Arabidopsis reference genome TAIR10 using STAR aligner^37^ with --outSAMtype BAM SortedByCoordinate --limitBAMsortRAM 16000000000 --outFilterType BySJout --alignSJoverhangMin 8 -- alignSJDBoverhangMin 1 parameters. Read counts for sense and antisense strands of each gene were calculated with --quantMode GeneCounts option of STAR command. To visualize mRNA coverage in IGV^38^ like in Fig. 1b, the “genomecov” function of BEDTools^39^ was used to make bedgraph files.

ChIP-seq data was processed as described in previously^11^. The ngs.plot program^40^ was used to make profiles for metaplots and heatmaps, which were then converted to log_2_ values and visualized in R and TreeView3 (https://bitbucket.org/TreeView3Dev/treeview3/src/master/). Sliding window analysis was performed using the “SlidingWindow” function of an R package evobiR (https://www.rdocumentation.org/packages/evobiR/versions/1.1/topics/SlidingWindow), with function: “mean”, window size: 150, and sliding size: 20.

For random forest analysis, every gene was divided into TSS region, TTS region, and gene body. The TSS (or TTS) region here was defined as 200 bp upstream and downstream from TSS (or TTS) combined, and the gene body as region spanning TSS to TTS. Short genes (< 400 bp) were omitted. Objective variable was defined as follows; genes with enriched ChIP-seq signal of FLD in the TTS regions, specifically RPM of FLAG-FLD – negative control > 20, are “FLD-bound” (n=3671), the others “unbound”. The number of the FLD-bound and unbound genes were balanced by randomly choosing 3671 from the unbound genes. Following predictor variables were generated by analyzing data of preceding works; H3K4me1, H3K4me2, H3K4me3, H3K9me2 and DNA methylation (mCG, mCHG, mCHH)^11^, H3K9 acetylation (H3K9ac) and H3K18ac^41^, H3.1 and H3.3^42^, H1 and H2A.Z^43^, H3K14ac^44^, LHP1 (Like-Heterochromatin Protein 1)^45^, H3K56ac, H3K36me3, and H3K27me3^46^, H3K23ac and H4K16ac^47^, H2B ubiquitination (H2Bub)^48^. The other features were from this study. Random forest was trained with genes on chromosome 1 – 4. Cross validation was performed with genes on chromosome 5. R package randomForest (https://www.rdocumentation.org/packages/randomForest/versions/4.6-14/topics/randomForest) was used with option ntree=1000. ROC curve was plotted using R package ROCR^49^.

For chrRNA-seq, read 1 of paired-end 150bp reads were trimmed for adaptors using Cutadapt^50^ with -a AGATCGGAAGAGCACACGTCTGAACTCCAGTCA option. Then 5’ poly-T bases derived from poly-A in 3’-ends of RNA were removed using Cutadapt with -g “XT{151}” --minimum-length 40 options. The trimmed reads were then mapped onto the TAIR10 reference genome using STAR aligner as described above for mRNA-seq. Downstream analyses were performed as described for mRNA-seq.

---

**Extended Data Fig. 1.**
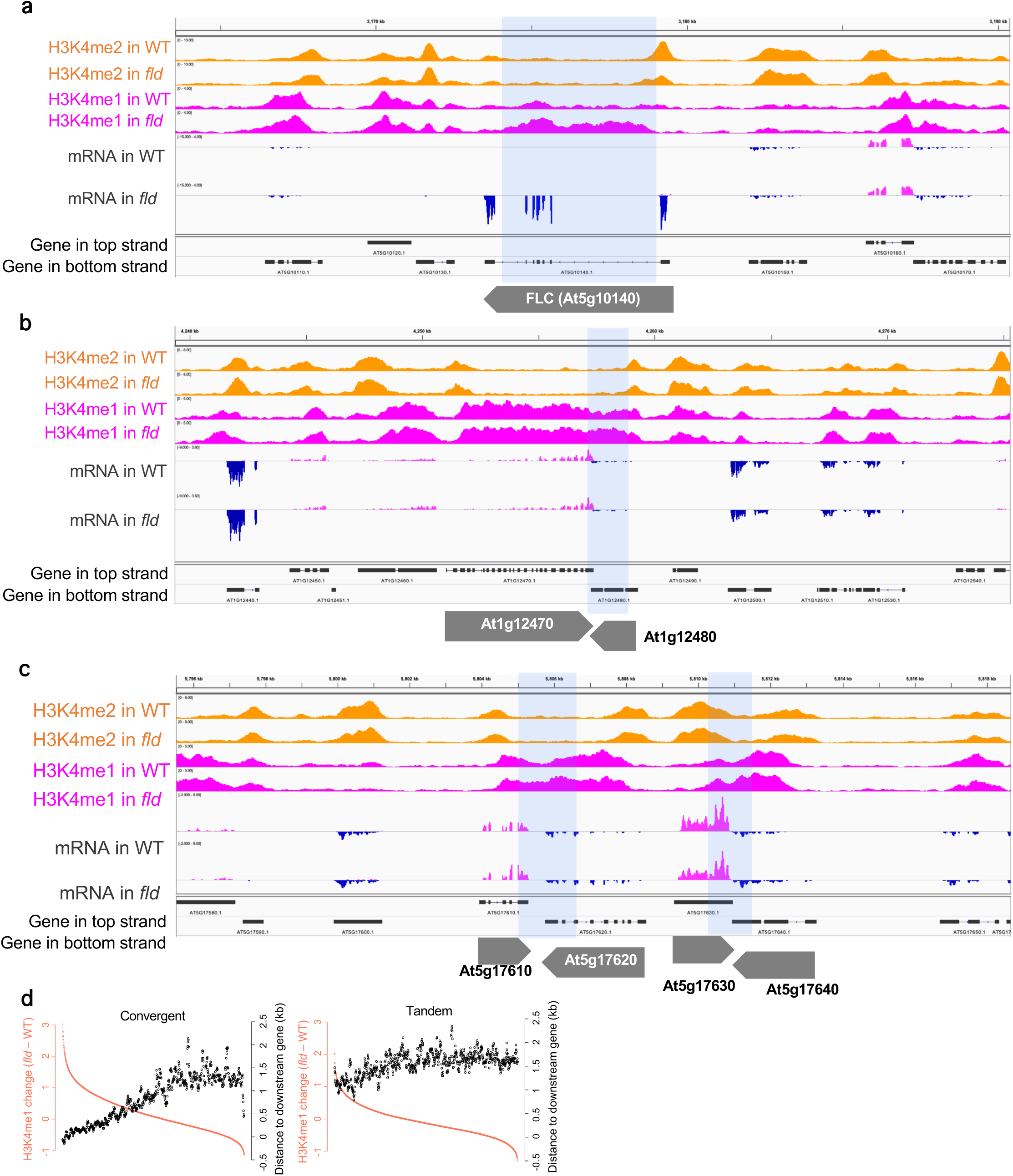
Effects of the *fld* mutation on H3K4me1 at the *FLC* locus and in convergent gene pairs. **a-c**, Patterns of H3K4me2, H3K4me1, and mRNA (top strand, magenta; bottom strand, dark blue) around the *FLC* locus (**a**, extended view of Fig. 1b) and pairs of convergent genes (**b, c**). Normalized coverage (per million mapped reads) is shown. Gene structures (box, exon; line, intron) in the top and bottom strands are shown. The shaded areas have increased H3K4me1 levels in *fld* compared to WT. **d**, Sliding window analysis (window size, 150; sliding size 20) showing the distance to downstream genes in relation to H3K4me1 changes between WT and *fld*. All genes were separated into convergent and tandem genes based on the orientation of downstream genes and were ranked and grouped by the differences in H3K4me1 levels between WT and *fld* (red plots and the left axis). The distances to downstream genes are shown by black plots and the right axis.

**Extended Data Fig. 2.**
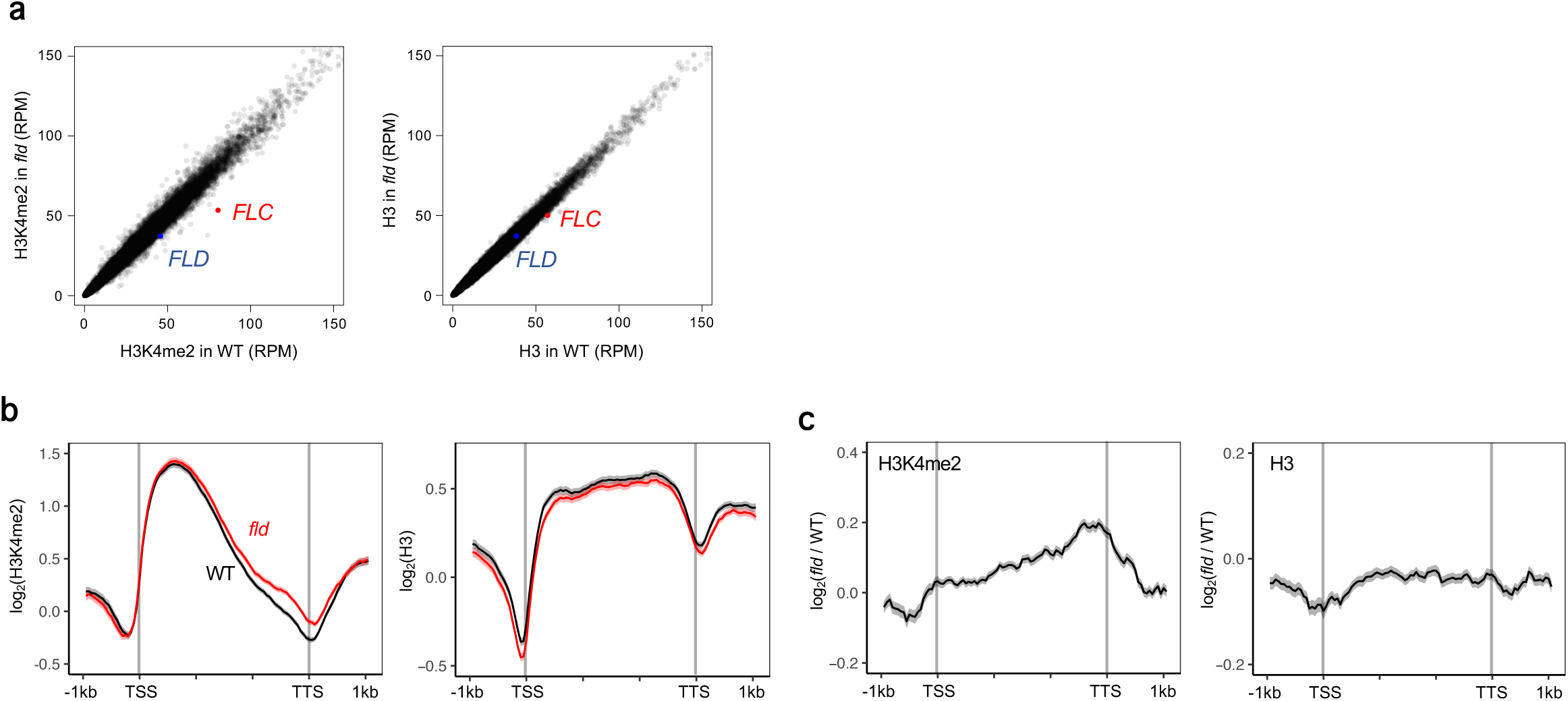
Effects of the *fld* mutation on H3 and H3K4me2 levels. **a**, Scatter plot comparing H3K4me2 (left) and H3 (right) levels between WT and *fld*. Each dot represents the RPM within each transcription unit (gene). **b**, Averaged profiles of H3K4me2 (left) and H3 (right) in WT and *fld.* **c**, H3K4me2 (left) and H3 (right) changes between WT and *fld* around genes with increased H3K4me1 in *fld* (1842 genes; same genes as Fig. 1c). The ribbons in **b** and **c** indicate the SEMs.

**Extended Data Fig. 3.**
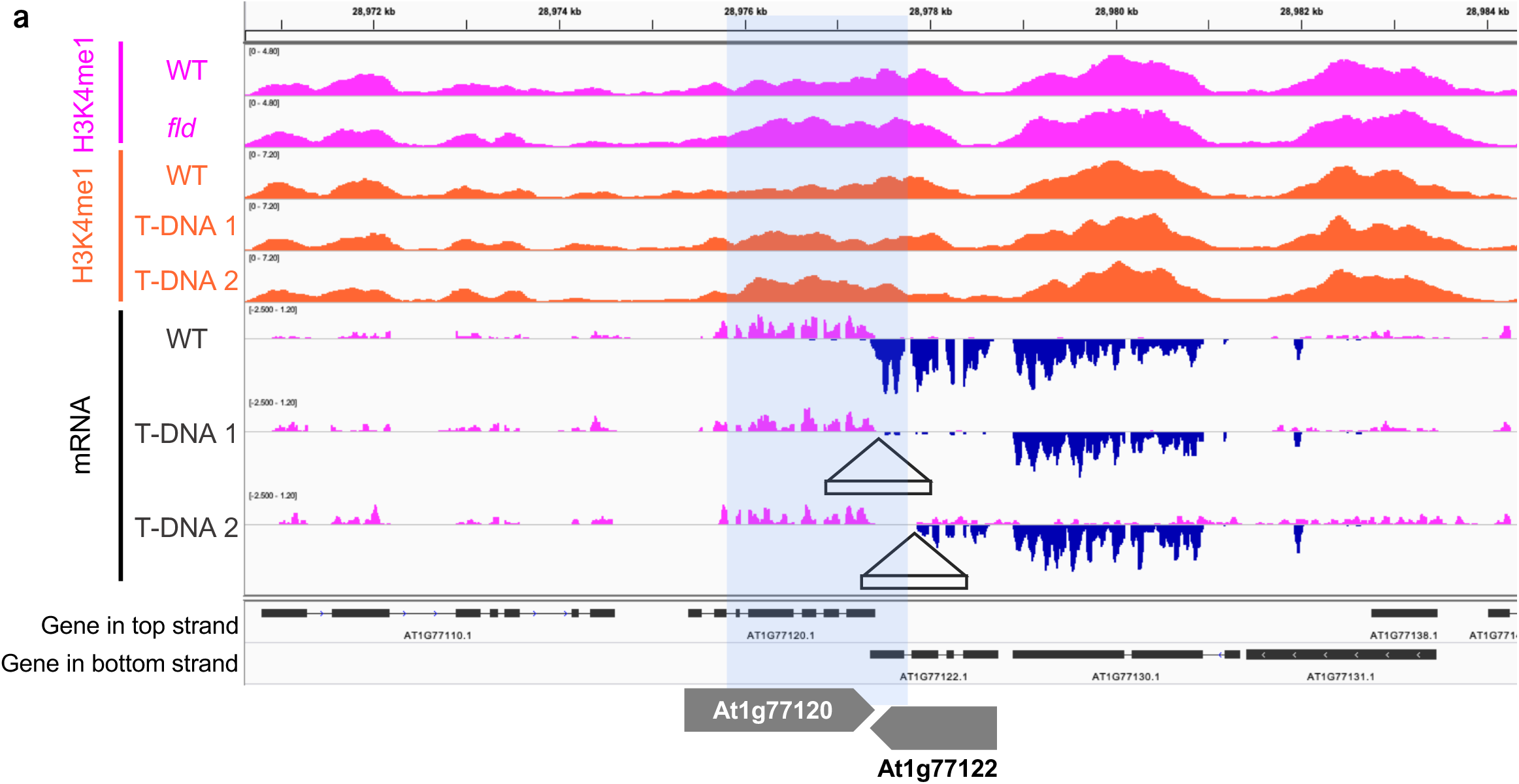
Eeffects of convergent transcription on H3K4me1. **a**, Patterns of H3K4me1 and mRNA in the At1g77120-At1g77122 pair in WT and two T-DNA insertion lines. The locations of T-DNA insertions are indicated by open triangles. The shaded areas have increased H3K4me1 levels in *fld* compared to WT.

**Extended Data Fig. 4.**
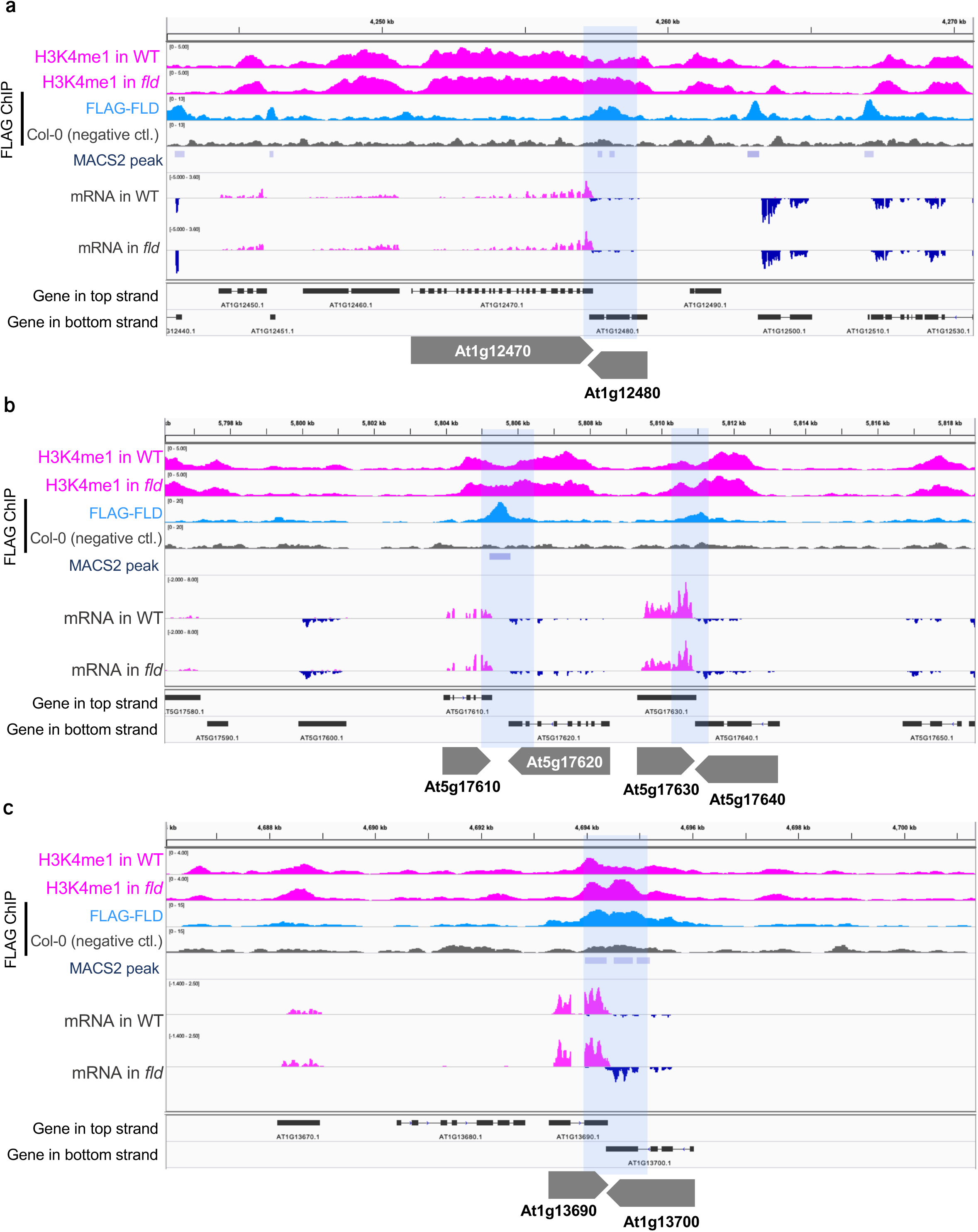
Localization pattern of the FLD protein. Patterns of H3K4me1, mRNA (top strand, magenta; bottom strand, dark blue), and FLD localization determined by ChIP-seq of FLAG-FLD. FLAG ChIP-seq of nontransgenic WT Col-0 plants was used as a negative control. The shaded areas have increased H3K4me1 levels in *fld* compared to WT.

**Extended Data Fig. 5.**
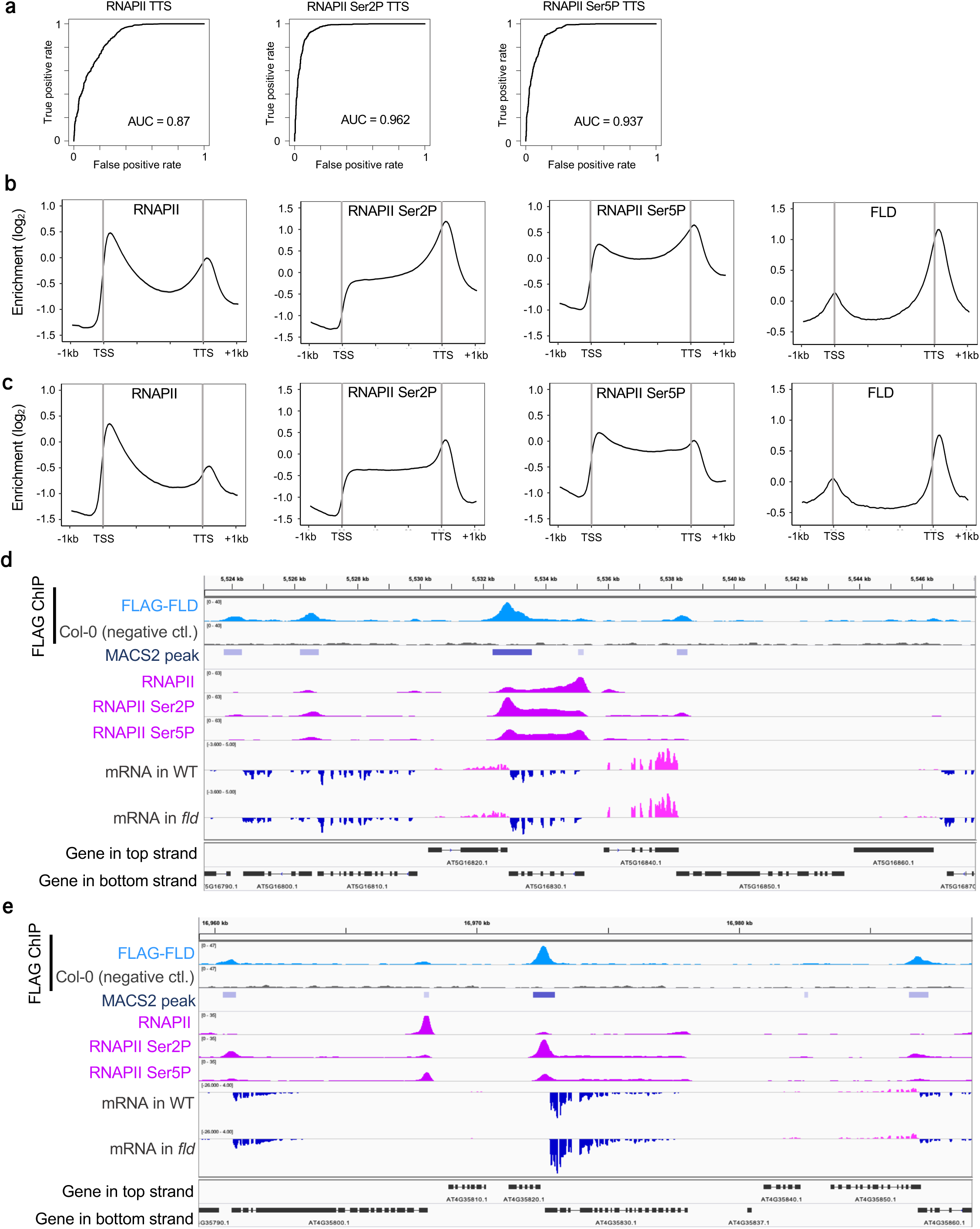
Colocalization of the FLD protein and phospho-RNAPII. **a**, Receiver-operator characteristic (ROC) curves showing FLC binding prediction accuracy of total RNAPII or phospho-RNAPII around TTS. A higher area under the ROC (AUC) value indicates higher accuracy as a predictor. **b, c**, Averaged profiles of RNAPII, RNAPII Ser5P, and RNAPII Ser2P and FLAG-FLD enrichment patterns around all convergent genes (**b**) and all tandem genes (**c**). **d, e**, RNAPII and FLAG-FLD profiles showing colocalization of FLAG-FLD and Ser2P and/or Ser5P of RNAPII.

**Extended Data Fig. 6.**
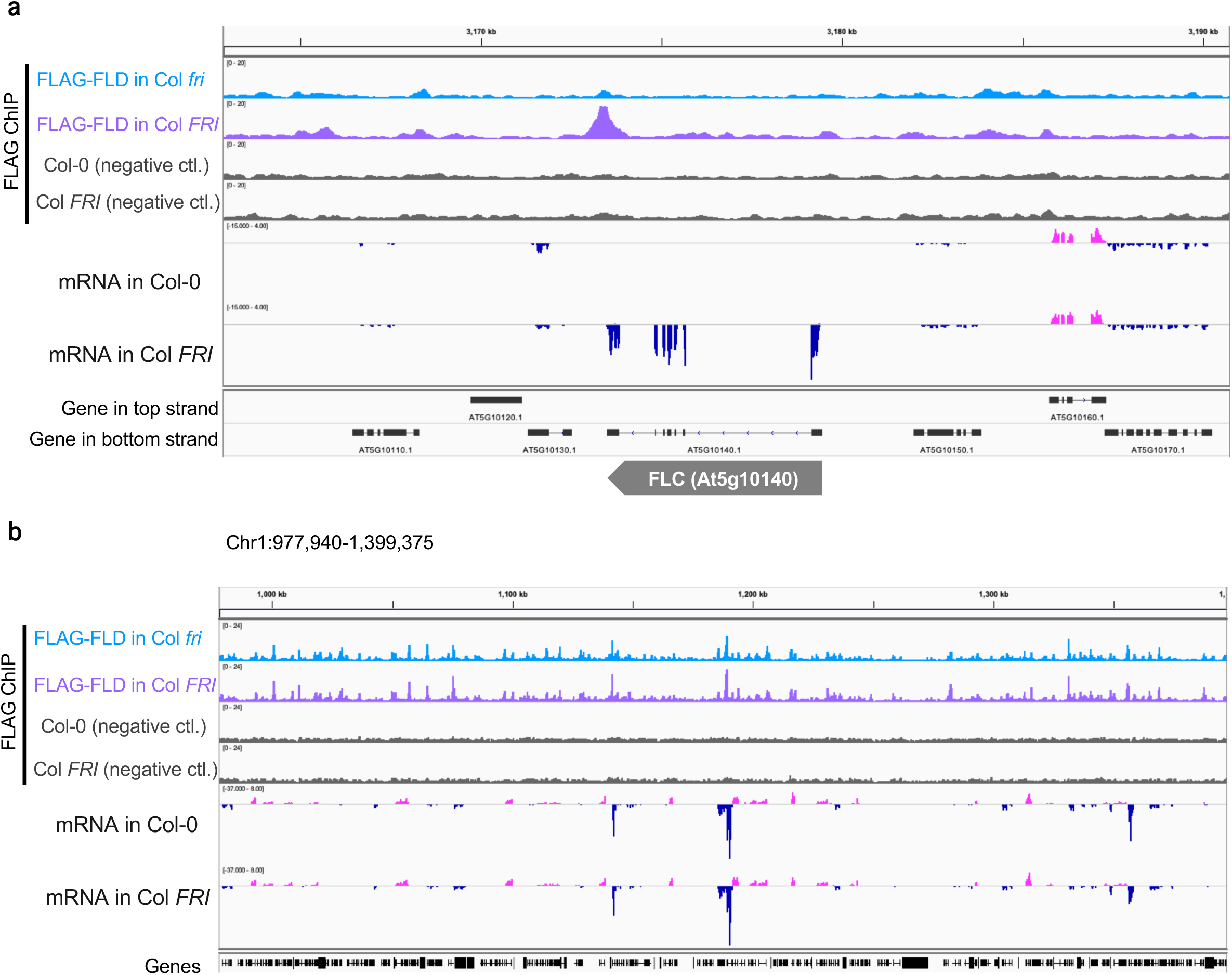
Effects of FRI-induced transcription on FLD localization. Patterns of FLD localization and mRNA (top strand, magenta; bottom strand, dark blue) comparing the *FLC* nonexpressor, Col *fri*, and the *FLC* expressor, Col *FRI*. FLAG ChIP-seq of nontransgenic Col-0 plants and Col *FRI* plants were used as negative controls. Extended views of Fig. 2d around the *FLC* locus (**a**) and the global pattern (**b**) are shown.

**Extended Data Fig. 7.**
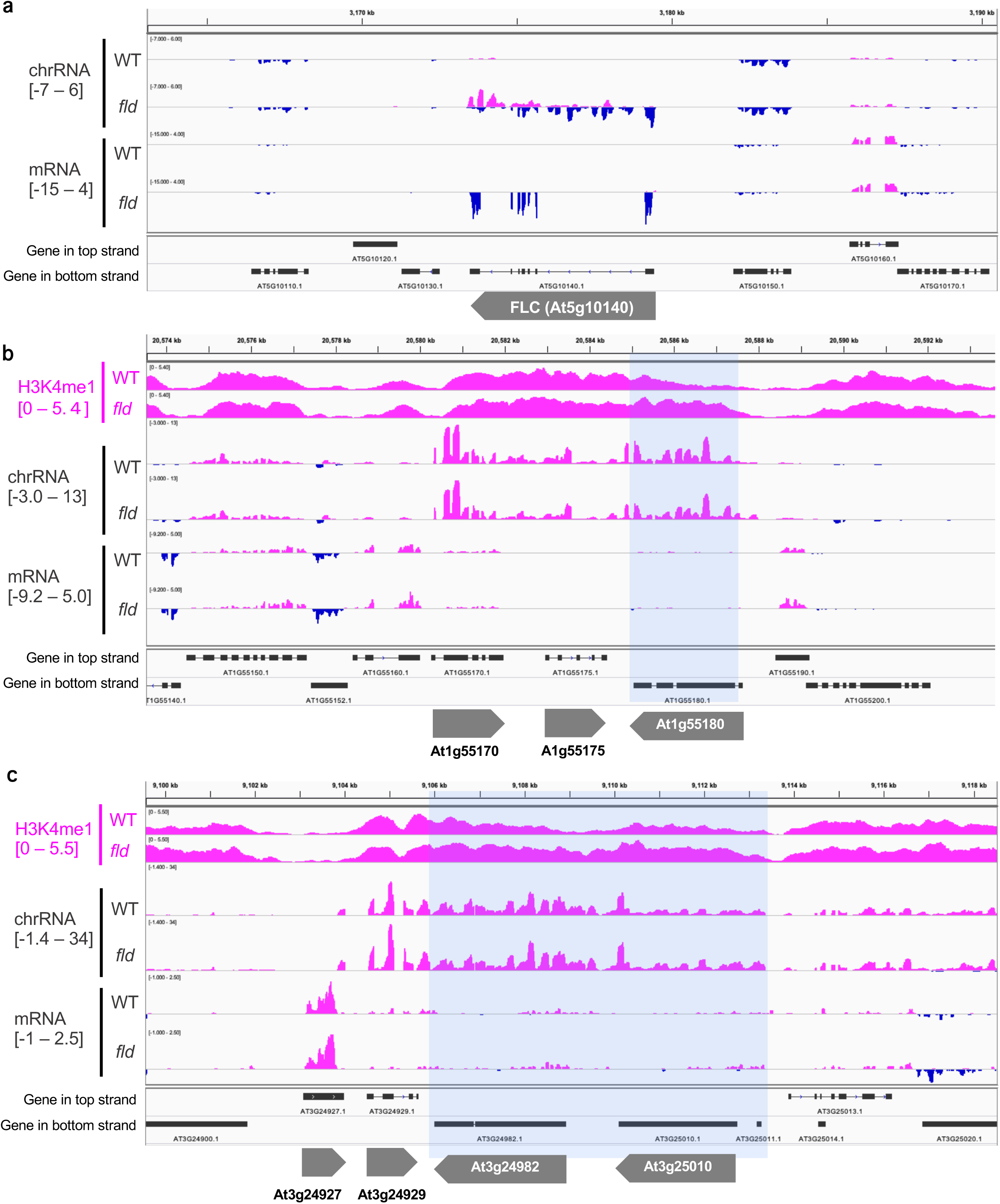
chrRNA and mRNA in WT and *fld*. **a**, Patterns of chrRNA and mRNA (top strand, magenta; bottom strand, dark blue) in WT and *fld* around the *FLC* locus. **b, c**, Patterns of H3K4me1, chrRNA, and mRNA around genes with highly increased H3K4me1 in *fld* compared to WT and with antisense chrRNA. The shaded areas have increased H3K4me1 levels in *fld* compared to WT.

**Extended Data Fig. 8.**
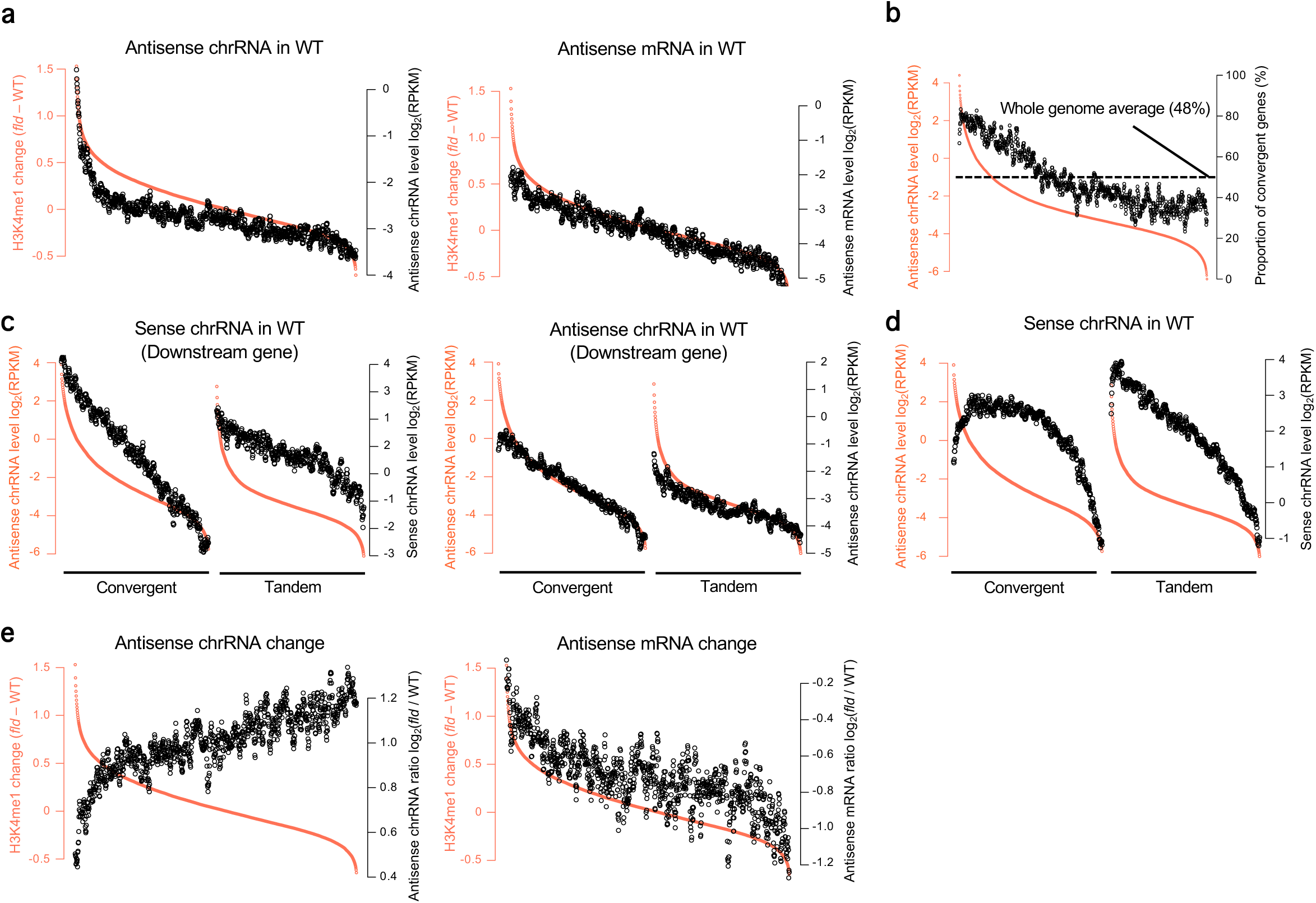
Antisense transcription, gene arrangement, and FLD-mediated H3K4me1 regulation. **a**, Comparison between chrRNA-seq (left, same as Fig. 3b) and mRNA-seq (right) for antisense transcripts. **b**, Sliding window analysis showing the proportion of convergent genes in relation to antisense chrRNA levels. All the genes were ranked and grouped by their antisense chrRNA level (red plots and the left axis). The proportion of convergent genes in each window is shown by black plots and the right axis. The dashed line indicates the proportion of convergent genes in the whole genome (48%). **c**, Sense (left) and antisense (right) transcripts of downstream genes in relation to antisense chrRNA levels. The genes were sorted and grouped like those in **b** but separately for convergent and tandem genes, and sense or antisense chrRNA levels (black plots) of downstream genes are plotted. **d**, Relationships between sense and antisense transcript levels. The genes were sorted and grouped like those in **c**, and sense chrRNA levels in WT are shown by black plots and the right axis. **e**, Effects of the *fld* mutation on antisense chrRNA (left) and antisense mRNA (right). All genes were grouped like those in **a**, and the log_2_ ratio of antisense RNA between WT and *fld* is shown by black plots and the right axis.

**Extended Data Fig. 9.**
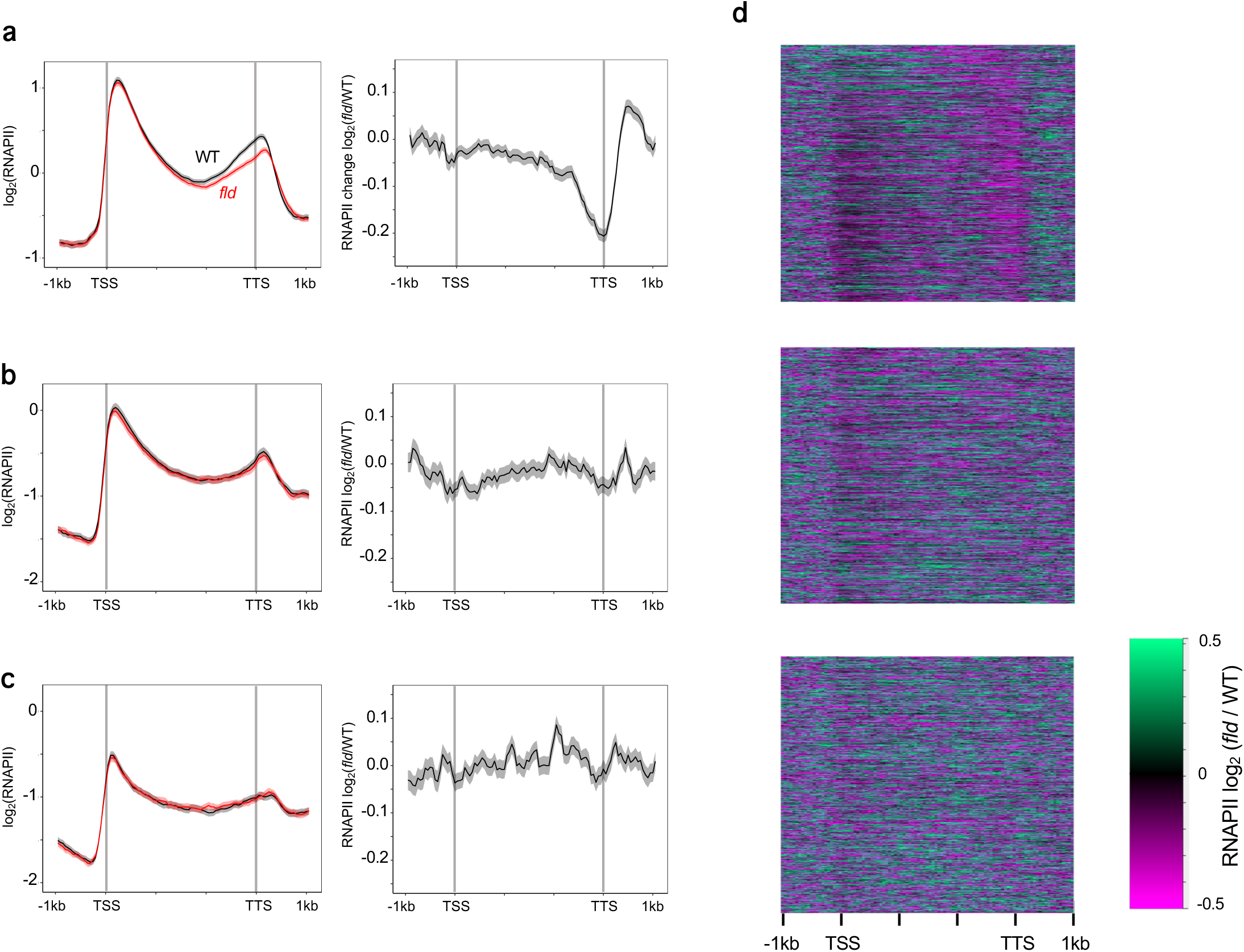
Effects of the *fld* mutation on the RNAPII pattern. **a-c**, Averaged profiles of RNAPII in WT and *fld* (left) and RNAPII changes (right) around genes categorized based on H3K4me1 increases in *fld* (top 1842 genes, **a**; middle 1842 genes, **b**; bottom 1842 genes, **c**). The ribbons indicate the SEMs. **d**, Heatmaps showing RNAPII changes between WT and *fld* around transcription units sorted by H3K4me1 increases in *fld*. The top, middle, and bottom panels correspond to the top 1842, middle 1842, and bottom 1842 genes, respectively.

**Extended Data Fig. 10.**
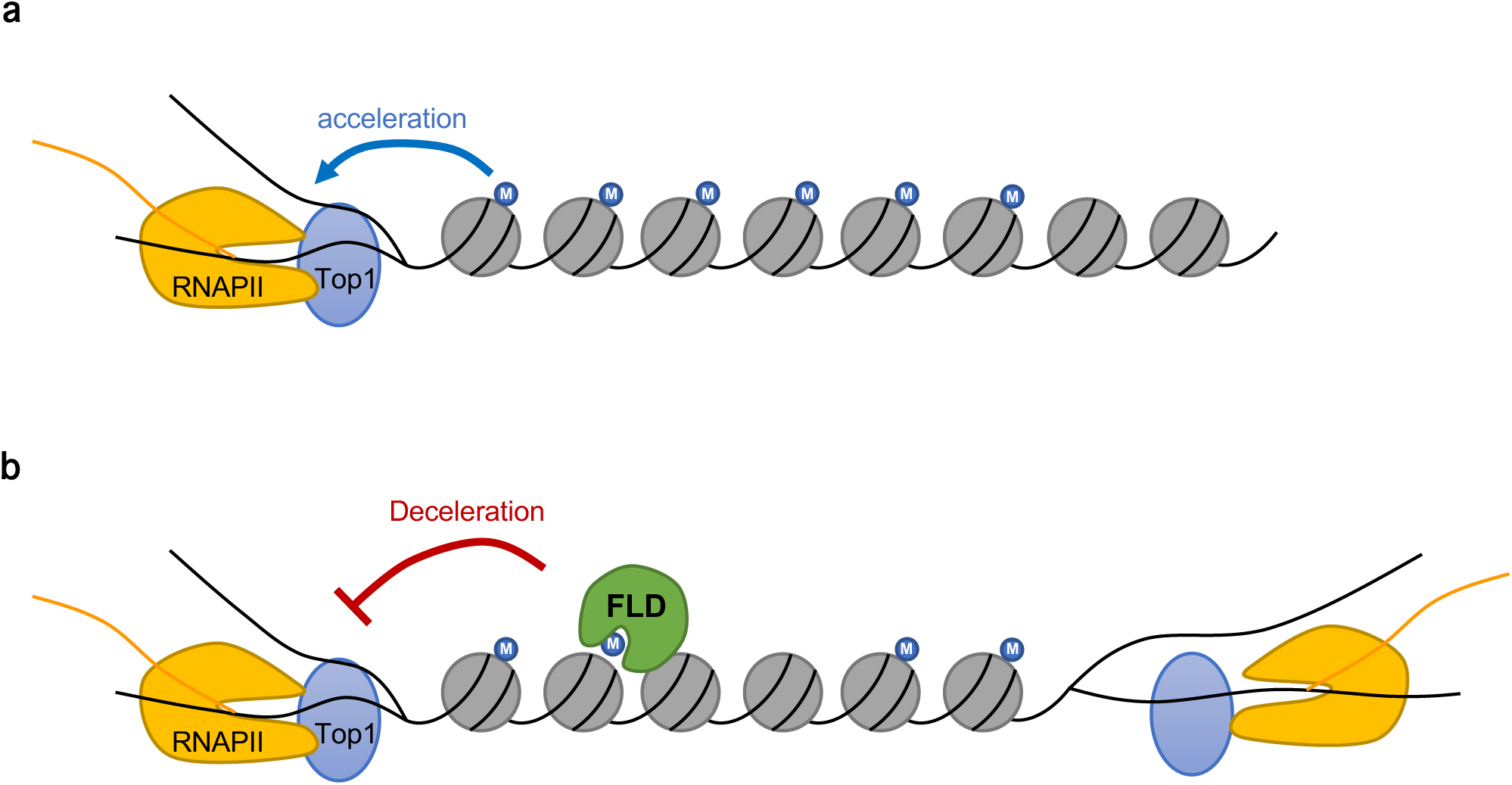
Model for the FLD-mediated regulation of convergent bidirectional transcription. **a**, RNAPII transcribes along the gene body with H3K4me1 (blue circle with “M”), with the help of Top1 to resolve supercoils. H3K4me1 increases the rate of transcription elongation possibly by activating Top1. **b**, At the regions of convergent overlapping transcription, FLD decelerates transcription elongation through removal of H3K4me1. In this scenario, we speculate that the activities of FLD and Top1 antagonize each other, which was supported by our genetic analyses.

